# Modified Long-Acting UCN2 Promotes Fat Mass Loss While Preserving Skeletal Muscle Mass and Function in Mouse Models of Pharmacological Weight Loss and Sarcopenic Obesity

**DOI:** 10.64898/2026.11.12.766237

**Authors:** Pablo Vidal, Patricia A.M. Baumgarten, Corey D. Holman, Jennifer R. Gamarra, Elizabeth R.M. Zunica, Dylan C. Seiler, Wagner S. Dantas, Analisa L. Taylor, Elizabeth C. Heintz, Emmanuel Akoto, Gabriel S. Gaidosh, Linsey Peed, Benjamin F. Miller, Mark R. Wade, Jill A. Willency, Steve M. Bauer, Samantha G.L. Keyser, Valentina Pirro, Joseph A. Baur, Paul M. Titchenell, John P. Kirwan, Christopher L. Axelrod, Joseph T. Brozinick

## Abstract

No approved pharmacotherapy simultaneously reduces excess adiposity and preserves skeletal muscle mass and function. Such an agent would have two distinct applications: improvement of the quality of pharmacological weight loss and treatment of sarcopenic obesity. Urocortin 2 (UCN2) and its receptor corticotropin-releasing hormone receptor 2 (CRHR2) are highly expressed in skeletal muscle, where they regulate hypertrophy and metabolism. Here we show that a modified long-acting acylated UCN2 peptide reduces fat mass while preserving muscle mass and enhancing function across distinct mouse models. Pair-feeding showed these effects were independent of reduced food intake. Combined with the GLP-1 receptor agonist semaglutide or the GIP/GLP-1/glucagon triple agonist retatrutide, UCN2 produced the greatest magnitude of fat loss while preserving lean and muscle mass and increasing *in vivo* force production, and this gain persisted 4 weeks after dosing stopped. UCN2 increased energy expenditure, prevented the adaptive fall in expenditure caused by reduced food intake, and promoted whole-body fatty acid oxidation. In a sarcopenic obese mouse model, UCN2 reduced fat mass while increasing muscle mass, muscle function, and myofibrillar protein synthesis. These findings identify UCN2-CRHR2 signaling as a strategy to uncouple fat loss from muscle loss, both as a standalone therapy for sarcopenic obesity and as a combination partner with incretins.

## Introduction

Skeletal muscle comprises 30-40% of adult body mass and is the primary site of insulin-mediated glucose uptake and insulin resistance in type 2 diabetes ^1^. Incretin therapies have transformed obesity treatment ^2–4^, but 25-40% of weight lost is lean mass ^2,3,5–8^. While some lean loss is expected during weight loss and may not impair function in healthy individuals ^9–12^, low muscle mass predicts mortality ^13^, and sarcopenic obesity, characterized by obesity in conjunction with low muscle function and mass, affects millions ^14^, yet no approved pharmacotherapy addresses adiposity and muscle deficit together ^8,15^. Activin type II receptor blockade preserves lean mass during GLP-1-induced weight loss ^12,15,16^, showing such combinations are feasible. Resistance exercise also protects muscle during GLP-1-induced weight loss ^17^, but long-term adherence is poor, motivating pharmacological alternatives. Urocortin 2 (UCN2) and its receptor CRHR2 are enriched in skeletal muscle, where UCN2 drives hypertrophy, enhances insulin sensitivity, and protects against diet-induced obesity ^18–23^. However, the effects of chronic pharmacological UCN2 treatment on muscle function remain unclear, and whether UCN2 can counteract incretin-induced lean mass loss is unknown. We therefore asked whether pharmacological UCN2 could 1) improve the efficacy and durability of incretin-induced weight loss, and 2) reverse the combined adiposity and muscle deficit of sarcopenic obesity

## Methods

### Animals

#### Mouse study 1: Diet-induced obesity mouse (retatrutide)

Experiments were approved by the Eli Lilly institutional animal care and use committee and were carried out in accordance with the National Research Council’s Guide for the Care and Use of Laboratory Animals. Male C57BL6/J DIO mouse model #380050 were purchased from Jackson Laboratories at 22 weeks of age. Mice were placed on a HFD at 6 weeks of age at Jackson laboratories and were already shipped as a diet-induced obesity model. Mice were single-housed in a temperature and humidity-controlled facility (22°C and 70% relative humidity) with a 12:12h light and dark cycle and *ad libitum* access to High Fat Diet (Research Diets D12492, 20% kcal from protein, 60% kcal from fat and 20% kcal from carbohydrates) and water, unless otherwise specified. After approximately 1 week of acclimation, mice were randomized and received intrascapular subcutaneous injections of vehicle (Tris-HCl), modified UCN2 (72 nmol/kg) every 3 days, or retatrutide (10 nmol/kg)^3^ every 3 days. Working solutions were prepared in 40 mM Tris pH 8 + 50 mg/mL of mannitol and 0.02% Polysorbate 80. In one study, mice received 6 doses and tissues were collected 48h after the last dose. Pairfed mice were staggered for one day, and each day at 9:00 am they received the average food intake of the UCN2 treated mice from the previous day. In the withdrawal study, mice received 5 doses over 14 days prior to discontinuation of treatment for 28 additional days. Mice that were placed on indirect calorimetry chambers for energy expenditure measurements were housed at 30°C for 1 week prior to the start of the study and for the duration of the study (14 days). Mice were sacrificed with CO_2_ followed by cervical dislocation, and tissues were harvested and snap frozen in liquid nitrogen and stored at -80°C until use.

#### Mouse study 2: Diet-induced obesity mouse (semaglutide)

Animal experiments were conducted in accordance with guidelines of the Institutional Animal Care and Use Committee at the University of Pennsylvania. Mice were maintained under 12-hour

light/dark cycles at ∼22°C. Diet-induced obese C57BL/6J males (#380050) were purchased from Jackson at 22 weeks of age and allowed to acclimate to single housing for one week prior to beginning the experiment. Mice had free access to a High-fat diet containing 60% kcal from fat (Research Diets #D12492i) for the duration of the experiment. Semaglutide (provided by Eli Lilly) or equal volume vehicle control (0.9% saline) were administered subcutaneously at 30nmol/kg ^24^ body weight daily for two weeks. Modified UCN2 (provided by Eli Lilly) or equal volume vehicle control (40 mM Tris pH 8.0, 50 mg/mL D-Mannitol, 0.02% Polysorbate 80) were administered subcutaneously at 10 mL/kg body weight (72 nmol/kg) every three days for two weeks. The final dose of UCN2 was given two days prior to tissue harvest. Mice were sacrificed with CO_2_ followed by cervical dislocation, and tissues were harvested and snap frozen in liquid nitrogen and stored at -80°C until use.

#### Mouse study 3: Sarcopenic obesity mouse model

All procedures involving animals were approved by the Institutional Animal Care and Use Committee at Pennington Biomedical Research Center. Male C57BL/6J mice (n = 28) were purchased from Jackson Laboratory and maintained on a standard chow diet (Purina 5001). After an acclimation period, at 20 months of age, mice were single-housed at 27–28 °C with a relative humidity of 50 ± 10% on a 12 h light:dark cycle, with ad libitum access to food and water. Animals were randomized to either a low-fat diet (LFD; 10% fat, 20% protein and 70% carbohydrates, D12450J, Research Diets) or a high-fat diet (HFD; 60% fat, 20% protein, and 20% carbohydrate, D12492, Research Diets) for 8 weeks. Subsequently, the HFD group was further randomized to receive intrascapular subcutaneous injections of either vehicle (Tris-HCl) or modified UCN2 (30 nmol/kg) every 3 days for a total of 12 doses over a 4-week period. Working solutions were prepared in 20 mM Tris-HCl. Mice were sacrificed with CO_2_ followed by cervical dislocation, and tissues were harvested and snap frozen in liquid nitrogen and stored at -80°C until use.

### UCN2 peptide

Modified UCN2 peptide was synthesized using established solid-phase peptide synthesis protocols, purified using reversed-phased chromatography and conjugated to a fatty acid.

### Whole-Body Composition

Fat and lean mass were assessed at multiple timepoints throughout the studies by nuclear magnetic resonance using the EchoMRI Body Composition Analysis system (EchoMRI, Houston, TX) or LF110 BCA Analyzer (Bruker Corporation, Billerica, MA, USA). Mice were restrained in opaque tubes to minimize stress and movement. Measurements were made in the ad libitum fed state.

### Indirect calorimetry

Whole-body energy metabolism was assessed using an open-circuit indirect calorimetry system (Comprehensive Laboratory Animal Monitoring System [CLAMS], Columbus Instruments, Columbus, OH, USA or Promethion CORE Metabolic Cages, Sable Systems International, North Las Vegas, NV, USA). Mice were maintained on a standard 12:12-h light–dark cycle at an ambient temperature of 22°C (Promethion) or 30°C (CLAMS) with *ad libitum* access to food (except pairfed group) and water throughout the monitoring period. Prior to data collection, mice were acclimated in the metabolic chambers for 3 days (Promethion) or 7 days (CLAMS) and then housed in metabolic chambers for 4 days (Promethion) or 14 days (CLAMS) for the duration of the study.

The O_2_ and CO_2_ sensors were calibrated prior to each experiment per the manufacturer’s specifications. Measurements were collected at regular intervals throughout the 12-h light/12-h dark cycle. Cumulative food intake and water consumption were monitored gravimetrically. Spontaneous physical activity was recorded as the number of infrared beam breaks along the X-Y axis (ambulatory movements, defined as consecutive beam breaks) and Z-axis (rearing activity). Concurrent fecal calorimetry (IKA C 3000) was also measured in the Promethion Core Metabolic Cage System. Caloric loss to feces was calculated using standard methods ^25^. Data were processed and analyzed using CalR2 ^26^.

### Grip strength

Grip strength was assessed in the sarcopenic obesity mouse model using a commercially available grip strength meter (Bioseb, Florida). Mice were held by the base of the tail and allowed to grasp the wire grid with all limbs. The experimenter gently pulled backward until the grip was released. Peak muscular force production of all four limbs was determined across three trials, and the average of the three trials was used for evaluation.

### Wire hang

Wire hang was assessed in the sarcopenic obesity mouse model using a wire grid. After allowing the animal to grasp the grid with all four limbs, the grid was gently inverted so that the mouse hung upside down. The latency to fall from the grid was recorded. A maximum cutoff time (150 sec) was applied to avoid excessive fatigue. Mice that fell prematurely were allowed one additional trial, and the average hanging time was used for analysis.

### Aurora 3 in 1 muscle force assay

Mice were anesthetized with isoflurane, and the leg of interest was shaved using hair clippers. Animals were then positioned supine on the heated platform of the Aurora 1300B Integrated 3-in-1 Whole Animal System (Aurora Scientific, Toronto). The shaved leg was stabilized with a pin at the knee to keep it straight and parallel to the ground and minor adjustments were made to ensure full contact with the foot plate. The foot was secured to the pedal with tape to prevent movement during the assay. To measure force production in the gastrocnemius, needle electrodes were inserted into the medial belly of the gastrocnemius muscle, piercing the tissue minimally to avoid damage. To measure force production in the tibialis anterior, surface electrodes were used. Individual stimulations (100 Hz, 0.2 ms) were applied while adjusting the “Stimulation Adjust” (range: 100 mA) to determine the minimal current required to elicit maximal force. After a 1-minute rest, the “Force–Frequency” protocol was initiated, consisting of ten 0.5-second stimulation trains at increasing frequencies (10, 20, 40, 60, 80, 100, 120, 140, 160, and 180 Hz) with rest intervals of 30 s between 10, 20, and 40 Hz stimulations and 60 s for the rest of the frequencies. Maximum force at each frequency was recorded and plotted to generate a force–frequency curve.

### Skeletal muscle protein synthesis

Mice were labelled with deuterium as previously described ^27,28^. Mice were switched to water bottles at 10% D_2_O (Sigma-Aldrich, D2O, 99%) for 7 days prior to tissue collection. Additionally, mice were injected intra peritoneally with 100% D_2_O at a dose volume of 20mL/kg twice, 7 days and 6 days prior to tissue collection. Tissues were collected and freeze clamped.

### Skeletal muscle protein degradation

D_3_-N, Methylhistidine was administered via oral gavage at 1mg/kg concentration with a dose volume of 5ml/kg 24h prior to tissue collection. Urine samples were collected at timepoints 0h and 24h, and plasma samples were obtained via tail bleed at timepoints 0h, 12h, 14h, 16h, 18h and 24h. Tissues were collected and freeze clamped at timepoint 24h.

### Mass spectrometry analyses

#### Myofibrillar Protein Fractional Synthesis Rate Method 1

A previously published method to analyze myofibrillar protein synthesis was used ^27–29^. Briefly, ∼30-50 mg of skeletal muscle tissues was homogenized in a bullet blender with isolation buffer (1:20; 100 mM KCl, 40 mM Tris–HCl, 10 mM Tris Base, 5 mM MgCl2, 1 mM EDTA, 1 mM ATP, pH = 7.6) containing phosphatase and protease inhibitors. After homogenization, subcellular fractions were isolated by differential centrifugation to obtain myofibrillar subcellular fraction, which was hydrolyzed and underwent cation exchange chromatography. We used gas chromatography-mass spectroscopy (7890A GC-Agilent 5975C MS, Agilent, Santa Clara, CA) to determine the deuterium enrichment of alanine in the myofibrillar fraction. Body water enrichment of D_2_O was measured from 50 μL of plasma that was prepared for analysis as described and run on a liquid water isotope analyzer (Los Gatos Research, Los Gatos, CA, USA). The newly synthesized fraction (f) of proteins was calculated from the enrichment of alanine bound in muscle proteins over the entire labelling period, divided by the true precursor enrichment (p), using plasma D_2_O enrichment with MIDA adjustment.

#### Myofibrillar Fractional Synthesis Rate Method 2

Snap-frozen muscle samples were pulverized using a stainless-steel Bessman tissue pulverizer kept chilled in dry ice. Samples were homogenized using an ice-cold homogenization buffer, containing 50 mM Tris-HCl (pH 7.5), 1 mM EDTA, a phosphatase inhibitor (PhosStop), and an EDTA-free protease inhibitor cocktail (Roche, Mississauga, ON, CA). Samples were then centrifuged at 14,000 x g for 20 min at 4°C, and the supernatant was removed. Collagen was precipitated with 0.3 M sodium hydroxide added to each pellet after incubation at 37°C for 30 minutes and centrifugation at 14,000 x g for 20 minutes at 4°C, leaving a myofibrillar-enriched supernatant fraction. A 20% trichloroacetic acid solution was added to the supernatant and incubated on ice for 20 minutes. Samples were centrifuged again, the supernatant discarded, and the pellet washed with a 70% ethanol solution. Proteins were then hydrolyzed by adding a 6 M hydrochloric acid (HCl) solution, followed by overnight incubation at 110°C. Samples were centrifuged at 14,000 x g for 5 minutes at 4°C and supernatant diluted to a final 0.1 M HCl concentration. Free amino acids were isolated using a Strata X-C cartridge (Phenomenex, CA). Free amino acids were eluted with 1% ammonium hydroxide in 20% acetonitrile, following cartridge equilibration in methanol and wash using first 0.1% formic acid in water and then 0.1% formic acid in acetonitrile. Eluant was dried using a Genevac EZ 2 4.0 (SP Scientific, PA) and reconstituted in a mixture of acetonitrile-methanol 1:1 v/v for analysis by liquid chromatography coupled to high resolution mass spectrometry (LC-HRMS, Orbitrap Exploris 480, Thermo Scientific). Chromatographic separation of underivatized alanine was achieved using a Waters Acquity BEH Amide column (100 mm × 2.1 mm, 2.5 µm particle size) held at +40 °C. Elution solvents were 10 mM ammonium formate adjusted with 0.1% formic acid (mobile phase A) and 0.1% formic acid in acetonitrile (mobile phase B). Plasma aliquots from the same animals were deproteinized using ice-cold acetonitrile-methanol 1:1 v/v and centrifuged at 14,000 x g for 15 min at 4°C. Plasma alanine measurements were conducted using the same LC-HRMS methodology used for the muscle samples. Enrichment of deuterium in body water was measured by gas chromatography coupled to mass spectrometry using a methodology described elsewhere ^30^. The calculation of the myofibrillar fractional synthesis rate (FSR) was based on the product-precursor equation^31,32^.

#### Myofibrillar Fractional Breakdown Rate

Frozen plasma samples were thawed in an ice bath. A 10-µL aliquot was extracted with 150 µL of pre-chilled acetonitrile-methanol (1:1 v/v). The samples were mixed thoroughly and incubated at −20 °C overnight. After centrifugation, the supernatant was further diluted in the extraction solvent mixture for LC-MS/MS analysis using an Agilent 1290 Infinity II UHPLC system (Agilent, Santa Clara, CA) coupled with a SCIEX 6500+ triple quadrupole mass spectrometry system equipped with an electrospray source (SCIEX, Framingham, MA). Data were acquired using scheduled multiple reaction monitoring mode with positive polarity. Chromatographic separation was achieved using a Waters Acquity BEH Amide column (100 mm × 2.1 mm, 2.5 µm particle size) held at +40 °C. Elution solvents were 10 mM ammonium formate adjusted with 0.1% formic acid (mobile phase A) and 0.1% formic acid in acetonitrile (mobile phase B). A 0-1 % 3MH-d_3_ (Cambridge Isotope Laboratories, MA) enrichment curve was prepared as a serial dilution in acetonitrile-methanol 1:1 v/v from a stock solution in water to calculate the enrichment in the experimental samples. The myofibrillar fractional protein degradation rate was calculated by determining the atom percent excess (APE) which was then log-transformed. The decay rate (k) was derived from the negative slope of the linear regression of the log-transformed APE values over time, as shown in Eq. 1

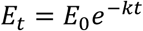

Where E_t_ is the enrichment of 3MH-d_3_ at time t following bolus tracer ingestion, and where E_0_ is the 3MH-d_3_ enrichment at the beginning of decay.

### Statistics

Comparisons between groups were analyzed using two-tailed Student’s t test, or two- or three-way analysis of variance (ANOVA; Vehicle vs UCN2 comparisons over time and/or condition) with Tukey’s HSD post hoc or Fisher’s LSD test applied where appropriate. ANCOVA test was performed for energy expenditure analyses with body mass-dependent parameters, based on recent consensus guidelines to preclinical indirect calorimetry experiments ^26^. Generalized linear models (GLM) with lean mass as a covariate were used for selected energy expenditure analyses. All statistical analyses were completed using GraphPad Prism software (version 10, GraphPad Software, La Jolla, CA). Significance was set at *p* < 0.05 and all data are presented as mean ± standard error of the mean (SEM).

## Main

We have generated a pharmacologically optimized modified acylated UCN2, hereafter referred to as UCN2, which dose-dependently reduces body weight, fat mass and food intake in diet-induced obese mice (Extended Data Fig. 1a–c). Previous data from our lab showed that UCN2 treatment decreased food intake ^18,19^. To understand the effects of UCN2 independent of reduced caloric intake, a pair-feeding experiment was performed (Fig. 1a). HFD fed mice were treated with Vehicle (*ad libitum*), UCN2 (*ad libitum*), or pair fed to the caloric equivalent of the UCN2 group (Fig. 1b,c). While pair-feeding also decreased body weight, UCN2 treatment resulted in a greater body weight reduction (Fig. 1d–f). UCN2 treatment was much more effective at decreasing fat mass compared to pair-feeding, while lean mass was dramatically decreased by only pair feeding but not UCN2 treatment (Fig. 1g). Pair-feeding did not increase muscle mass, force production or protein synthesis as UCN2 did (Fig. 1h–k). Thus the effects of UCN2 on body composition, muscle mass and function are not simply a consequence of reduced caloric intake. To assess energy expenditure, mice were housed at thermoneutrality (30°C) in CLAMS for 1 week of acclimatization prior to 2 weeks of treatment. The effects of UCN2 in body weight and composition were also recapitulated at thermoneutrality (Extended Data Fig. 2a–d). UCN2 treated mice had increased energy expenditure when compared to HFD Vehicle analyzed by ANCOVA, and UCN2 prevented the decrease in energy expenditure seen with decreased food intake, when compared to pairfed mice (Fig. 1l,m; Extended Data Fig. 2f–i). UCN2 treatment increased whole-body fatty acid oxidation that was constant throughout the treatment intervention, while the pairfed group showed a more variable pattern with a daily spike in RER when mice were given their caloric-matched food (Fig. 1n; Extended Data Fig. 2e). These data show that UCN2 treatment decreases food intake and increases energy expenditure, which explains the greater magnitude of weight loss when compared to just reduced food intake.

**Fig. 1.**
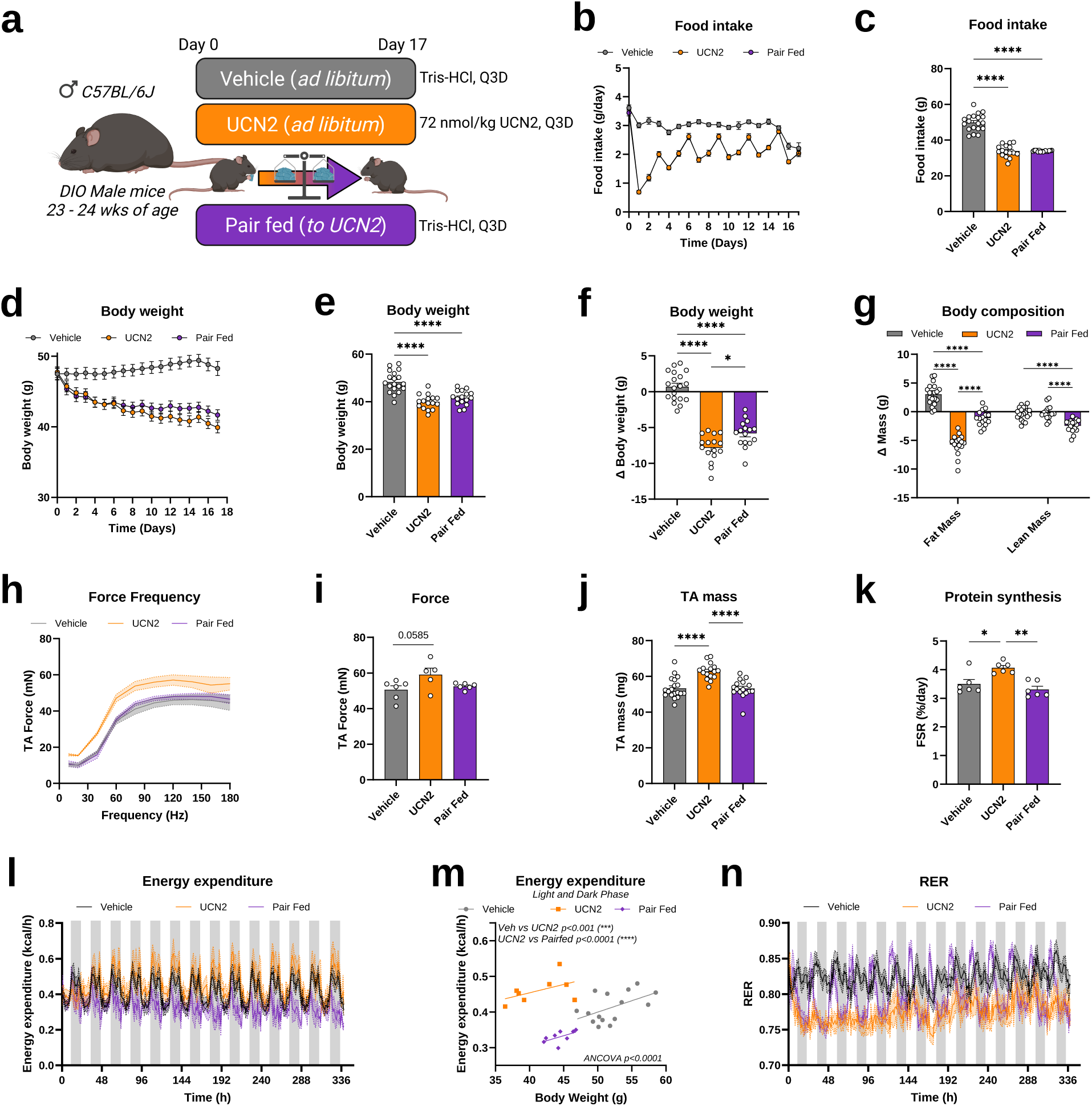
UCN2 preserves muscle mass and force production and increases energy expenditure independently of reduced food intake in obese mice. **a**, Schematic of the pair-feeding study. Young (24-week-old) diet-induced obese (DIO) male C57BL/6J mice were treated with vehicle (Tris-HCl, ad libitum), UCN2 (ad libitum) or pairfed to the caloric intake of the UCN2 group. Fig. 1b-k corresponds to mice housed at Room Temperature (20-22°C), and Fig 1l-n to mice housed at thermoneutrality (30°C) b, Daily food intake over the treatment period. c, Cumulative food intake. d, Body weight over the treatment period. e, Body weight at the end of treatment. f, Change in body weight. g, Change in fat mass and lean mass. h, In vivo TA force– frequency relationship. i, TA force production. j, TA muscle mass. k, Myofibrillar fractional synthesis rate (FSR). **l**, Energy expenditure over time by indirect calorimetry. **m**, ANCOVA of energy expenditure against body weight across the light and dark phases. **n**, Respiratory exchange ratio (RER) over time. Groups are vehicle, UCN2 and pairfed. Data are mean ± s.e.m.; n = 8-22 mice per group. Energy expenditure was analyzed by ANCOVA with body mass as a covariate; remaining comparisons used one- or two-way ANOVA with Tukey’s HSD post hoc test. *P < 0.05, **P < 0.01, ***P < 0.001, ****P < 0.0001.

Current incretin-based therapies for obesity cause a significant loss of fat and lean mass ^2–4^. To determine if that reduction of lean mass impacts muscle mass and function, and if UCN2 could preserve skeletal muscle during this pharmacological weight loss, young (24 weeks old) C57BL/6J DIO male mice were treated with vehicle, UCN2, semaglutide (sema) or the combination of UCN2 and sema (UCN2+Sema) (Fig. 2a). After the treatment period, mice treated with UCN2+Sema lost a greater amount of body weight compared to sema alone (Fig. 2b,c; Extended Data Fig. 3a,d,e). Body composition analysis revealed that mice treated with UCN2 or semaglutide lost a significant amount of fat mass, and the combination treatment resulted in greater fat mass loss without changes in lean mass (Fig. 2d; Extended Data Fig. 3b,f). UCN2-treated mice had a significant increase in TA weight compared to vehicle-treated mice (Fig. 2e), and UCN2 treatment prevented the sema-induced decreased in gastrocnemius weight (Extended Data Fig. 3c). Indirect calorimetry over the initial 4 days of treatment (Sable Systems Promethion) showed that both UCN2 and UCN2+Sema treated mice had a significant increase in energy expenditure compared to vehicle mice when analyzed by ANCOVA for total body mass (Fig. 2h,i; Extended Data Fig. 3g,j–l). Mice treated with UCN2, sema, or UCN2+Sema had a decrease in RER compared to control mice (Fig. 2j,k), which was significant even in the first 24h of treatment, replicating the adaptations seen at thermoneutrality (Extended Data Fig. 3m,n). Further analyses in the Promethion determined that sema alone or in combination with UCN2 significantly decreased food intake, body mass, and energy balance compared to vehicle mice looking at the initial 4 days of treatment (Fig. 2f,g,l,m; Extended Data Fig. 3d,e,h,i,o,p). Fecal calorimetry showed that mice treated with UCN2, semaglutide, or the combination had a significant decrease in the calories lost to feces, but this did not significantly alter energy balance conclusions when accounted for (Fig. 2n,o).

**Fig. 2.**
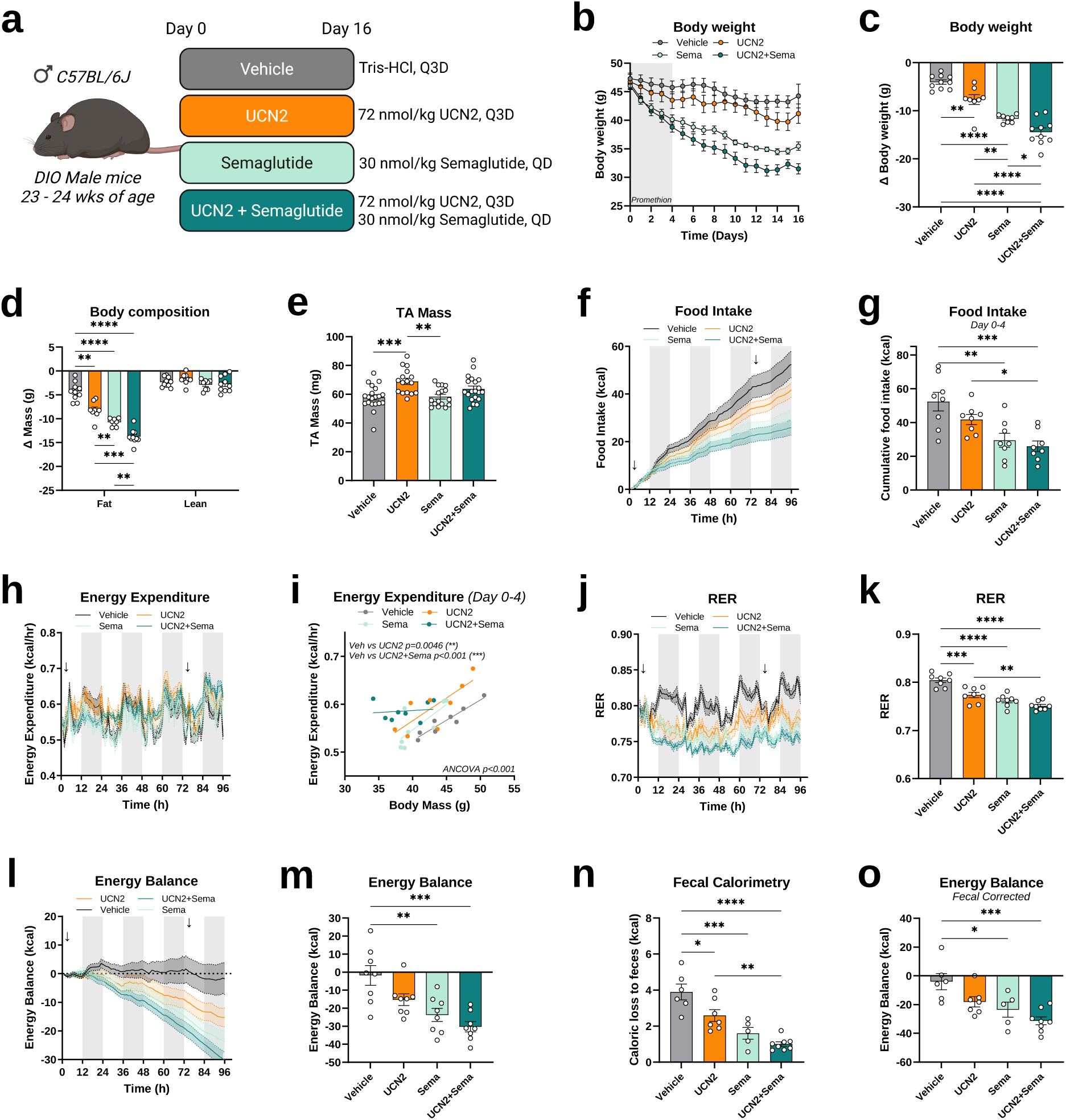
Combining UCN2 with semaglutide enhances fat mass loss and preserves muscle mass by increasing energy expenditure and fat oxidation. **a**, Schematic of the study design. Young (24-week-old) diet-induced obese (DIO) male C57BL/6J mice were treated with vehicle, UCN2, semaglutide (sema) or UCN2 plus sema, with the first 4 days of treatment recorded in the Promethion indirect-calorimetry system. **b**, Body weight over the treatment period. **c**, Change in body weight. **d**, Change in fat mass and lean mass. **e**, TA muscle mass. **f**, Food intake over the first 96 h of treatment. **g**, Cumulative food intake over the first 96h of treatment. **h**, Energy expenditure over the first 96 h of treatment. **i**, Average energy expenditure over the first 96 hrs of treatment versus total body mass with ANCOVA. **j**, Respiratory exchange ratio (RER) over the first 96 h of treatment. **k**, Mean RER. **l**, Energy balance over the first 96 h of treatment. **m**, Energy balance. **n**, Caloric loss to feces measured by bomb calorimetry. **o**, Energy balance corrected for fecal caloric loss. Groups are vehicle, UCN2, semaglutide (sema) and UCN2 + Sema. Data are mean ± s.e.m.; n = 8-10 mice per group. Energy expenditure was analyzed by ANCOVA with body mass as a covariate; remaining comparisons used one- or two-way ANOVA with Tukey’s HSD post hoc test. *P < 0.05, **P < 0.01, ***P < 0.001, ****P < 0.0001.

Retatrutide (reta) is a triple GLP1, GIP and Glucagon receptor agonist ^33–35^ that reduces body weight by up to 30.3% in humans (TRIUMPH-1). To determine if UCN2 could preserve muscle mass during profound weight loss, young (24 weeks old) DIO male C57BL/6J mice were treated with vehicle, UCN2, reta or the combination of UCN2 and reta (UCN2+Reta) (Fig. 3a). Reta alone or with UCN2 markedly reduced daily and cumulative food intake compared to HFD vehicle mice, and UCN2 alone reduced cumulative intake to a lesser extent (Fig. 3b,c; Extended Data Fig. 4a). Water intake did not differ among groups (Extended Data Fig. 4b–d). Following a similar pattern to food intake, UCN2 and reta reduced body weight by 17.34% and 31.9% respectively, and UCN2+Reta resulted in the greatest body weight loss over the 17-day treatment period (37.8%) (Fig. 3d,e; Extended Data Fig. 4e). Body composition analyses after 2 weeks of treatment showed that all treatment groups significantly reduced fat mass compared to HFD vehicle, which gained 18% fat mass during the study duration, with UCN2 reducing fat mass by 34.15%, reta by 55.90% and UCN2+Reta by 78.95% (Fig. 3f; Extended Data Fig. 4f). Reta treated groups had a significant loss of lean mass, while UCN2 alone preserved lean mass. UCN2 treatment decreased liver weight by ∼33%, while reta treatment alone or the combination decreased liver weight ∼50% (Extended Data Fig. 4g). The greater reduction of body weight seen with the combination of UCN2 and reta is explained by a greater reduction of fat mass without a further decrease in lean mass when compared to reta alone (Fig. 3f).

**Fig. 3.**
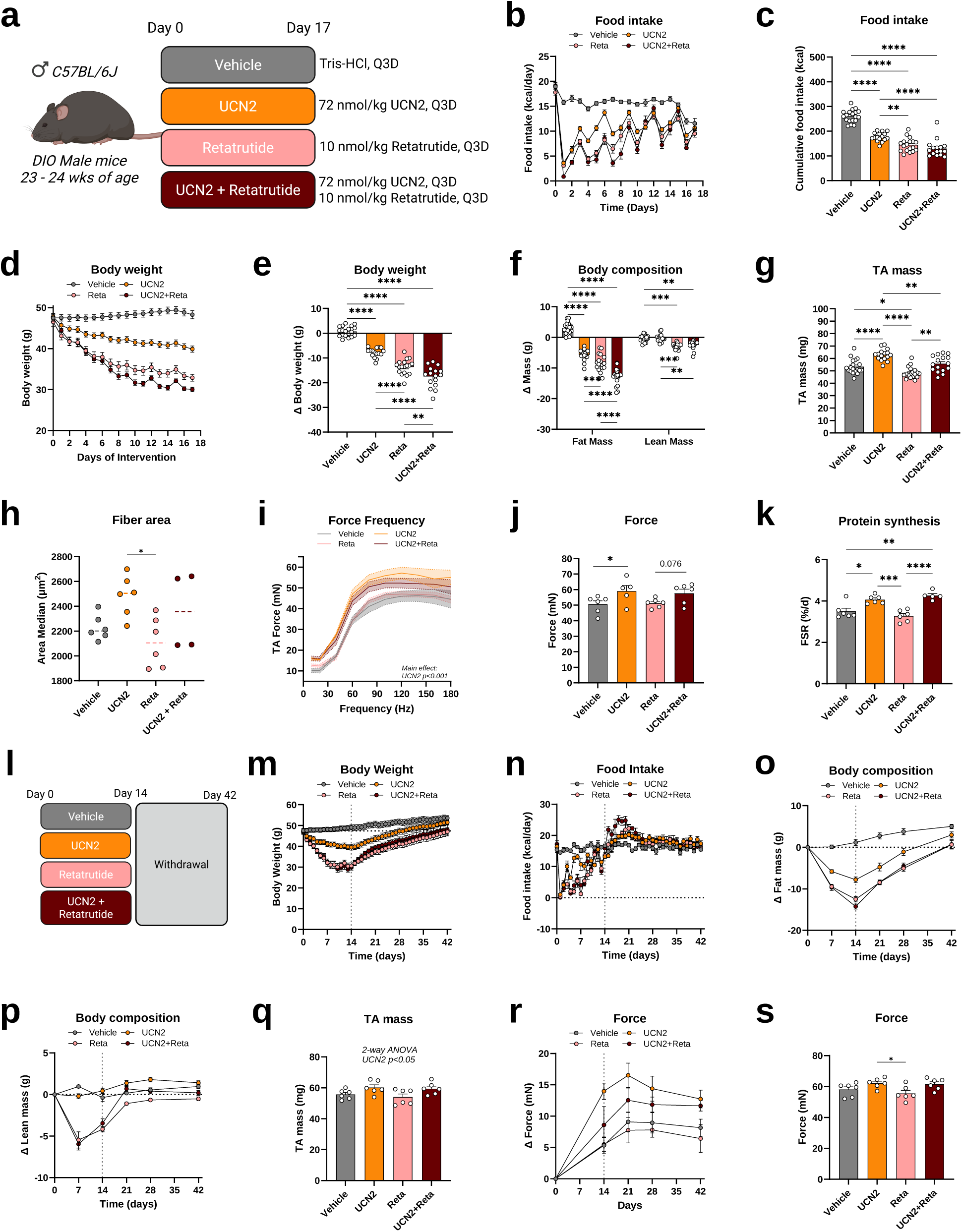
Combining UCN2 with retatrutide enhances fat mass loss and increases in vivo force production. **a**, Schematic of the study design. Young (24-week-old) DIO male C57BL/6J mice were treated with vehicle, UCN2, retatrutide (reta) or UCN2 plus reta for 17 days. **b**, Daily food intake. **c**, Cumulative food intake. **d**, Body weight over the intervention. **e**, Change in body weight. **f**, Change in fat mass and lean mass. **g**, TA muscle mass. **h**, Median TA fiber cross-sectional area. **i**, *In vivo* TA force–frequency relationship. **j**, TA tetanic force production. **k**, Myofibrillar fractional synthesis rate (FSR). **l**, Schematic of the withdrawal study. Mice were treated for 2 weeks and then followed for a further 4 weeks after cessation of treatment. **m**, Body weight during treatment and withdrawal. **n**, Food intake during treatment and withdrawal. **o**, Change in fat mass during treatment and withdrawal. **p**, Change in lean mass during treatment and withdrawal. **q**, TA muscle mass at the end of the withdrawal period. **r**, Change in in vivo TA force during treatment and withdrawal. **s**, TA force production at the end of the withdrawal period. Vertical dotted line indicates withdrawal start. Groups are vehicle, UCN2, retatrutide (reta) and UCN2 + Reta (UCN2+Reta). Data are mean ± s.e.m.; n = 6-22 mice per group. Mice from Vehicle and UCN2 groups are the same as shown in Fig 1 b-k. All groups were run together. Comparisons used one-, two- or three-way ANOVA with Tukey’s HSD post hoc test as appropriate. *P < 0.05, **P < 0.01, ***P < 0.001, ****P < 0.0001.

To assess the functional consequences of these changes in lean mass, we measured *in vivo* contractile force of the tibialis anterior (TA) muscle across a range of stimulation frequencies. UCN2 treated mice had a significant upward shift in the force frequency curve compared to vehicle mice (Fig. 3i). Despite the decrease in lean mass, there was not a decrease in muscle force production with reta, while the combination group also maintained force production above reta alone (Fig. 3i). These results were also found when measuring force production by the gastrocnemius (Extended Data Fig. 4m). Tetanus-induced force production in TA muscle was significantly increased in UCN2 treated mice compared to HFD vehicle, and UCN2 plus reta produced a non-significant increase when compared to reta alone (Fig. 3j). TA mass was increased 17.74% by UCN2 and reduced 9.18% by reta alone; the combination preserved TA mass relative to reta alone (+15.02%, and +4.46% vs. vehicle) (Fig. 3g). This increase in muscle weight also resulted in increased fiber size area in the UCN2 treated groups (Fig. 3h; Extended Data Fig. 4i–k). When force production was normalized to TA weight, there were no differences between groups, suggesting the increase in force production is mostly caused by increased muscle hypertrophy (Extended Data Fig. 4l). Consistent with the sarcopenic obesity model, UCN2 alone and the combination of UCN2 plus reta significantly increased the fractional synthesis rate compared to HFD vehicle and reta alone (Fig. 3k). While there were no significant differences in protein turnover compared to the HFD vehicle, UCN2 treated mice had lower protein turnover when compared to the retatrutide treated mice (Extended Data Fig. 4h).

While incretin treatment is very effective at causing weight loss, stopping treatment may result in a reversal of the improvements in body weight and composition. To understand if the positive effects to muscle function follow the same pattern, we dosed a cohort of mice for 2 weeks, which were then followed for 4 more weeks after cessation of treatment (Fig. 3l). Treatment cessation caused food intake to briefly increase prior to returning to levels of the vehicle group (Fig. 3n; Extended Data Fig. 5a,b). Body weight and body composition started to increase as soon as treatment was stopped and returned to almost baseline values by 4 weeks in all groups (Fig. 3m,o,p; Extended Data Fig. 5c,e–h). However, the increase in muscle function induced by UCN2, alone or in combination with retatrutide, was maintained up to 4 weeks after treatment was stopped (Fig. 3r,s; Extended Data Fig. 5d). While there were no differences by multiple comparisons, there was an overall 2-way ANOVA effect for UCN2 to increase muscle mass by 9% (Fig. 3q). These data highlight that UCN2 not only increases muscle mass and function, but these adaptations are preserved after treatment is stopped and the improvement in body composition is lost.

To determine if UCN2 could attenuate sarcopenic obesity, 20-month-old male C57BL/6J mice were fed a low fat diet (LFD) or a high fat diet (HFD) for 8 weeks, followed by 4 weeks of treatment with vehicle or UCN2 (Fig. 4a). UCN2 reduced daily and cumulative food intake compared to HFD vehicle mice (Fig. 4b,c), and attenuated weight gain over the 4-week treatment period, although total weight gain remained higher than in LFD vehicle mice (Fig. 4d,e). Body composition after 4 weeks of treatment showed that UCN2 treated mice had a significant reduction in fat mass compared to HFD vehicle mice (Fig. 4f,g). UCN2 treated mice gained lean mass over the treatment period, and this change in lean mass was significantly greater than HFD vehicle mice (Fig. 4h,i).

**Fig. 4.**
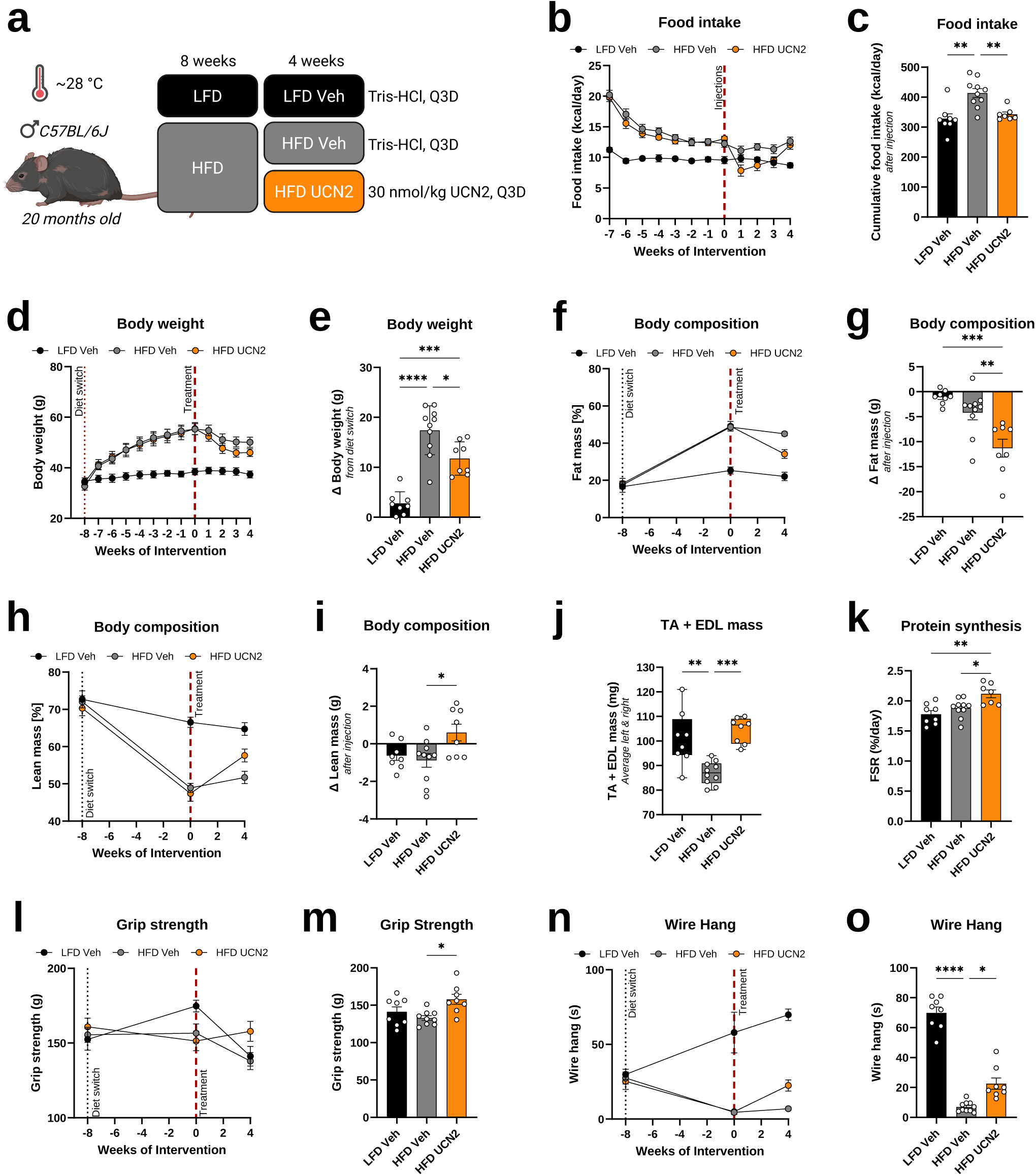
UCN2 treatment reduces high-fat-diet-induced weight gain and preserves muscle mass and function in mice with sarcopenic obesity. a, Schematic of the study design. Twenty-month-old male C57BL/6J mice were fed a low-fat diet (LFD) or a high-fat diet (HFD) for 8 weeks and then treated for 4 weeks with vehicle (Tris-HCl) or UCN2. b, Daily food intake over the intervention. **c**, Cumulative food intake over the treatment period. **d**, Body weight from diet switch through the end of treatment. **e**, Change in body weight from the diet switch. **f**, Fat mass (% of body mass) over the intervention. **g**, Change in fat mass over the treatment period. **h**, Lean mass (% of body mass) over the intervention. **i**, Change in lean mass over the treatment period. **j**, Combined tibialis anterior (TA) and extensor digitorum longus (EDL) mass (average of left and right limbs) at the end of the study. **k**, Myofibrillar fractional synthesis rate (FSR). **l**, Grip strength over the intervention. **m**, Grip strength after 4 weeks of treatment. **n**, Wire-hang duration over the intervention. **o**, Wire-hang duration after 4 weeks of treatment. Groups are LFD vehicle, HFD vehicle and HFD UCN2. Data are mean ± s.e.m.; n = 8-10 mice per group. Time-course data were analyzed by two-way ANOVA with Tukey’s HSD post hoc test and endpoint data by one-way ANOVA with Fisher’s LSD test. *P < 0.05, **P < 0.01, ***P < 0.001, ****P < 0.0001.

To assess skeletal muscle function, we measured grip strength and wire hang duration after 4 weeks of treatment. UCN2 treated mice had improved grip strength compared to HFD vehicle mice after the treatment period (Fig. 4l,m). Wire hang time progressively declined in mice that were fed a HFD, but UCN2 treatment increased their wire hang performance, although it was still lower than the Lean controls (Fig. 4n,o). To test whether this reflected hypertrophy, muscles were collected and weighed. TA + EDL mass was significantly increased in UCN2 treated mice compared to HFD vehicle, and was similar to LFD vehicle, preserving the mass lost on HFD (Fig. 4j). UCN2 also significantly increased the in vivo myofibrillar fractional synthesis rate (FSR) compared to HFD and LFD vehicle mice (Fig. 4k). These data demonstrate that UCN2 preserves skeletal muscle mass and function in aged, sarcopenic obese mice while promoting fat mass loss.

Sarcopenic obesity is associated with impaired mobility, metabolic dysfunction, and increased mortality ^8,14,36,37^. In aged HFD-fed mice, UCN2 reduced fat mass while restoring skeletal muscle mass to that of LFD vehicle controls and improving grip strength and wire hang performance. This dual action distinguishes UCN2 from current anti-obesity therapies, which reduce fat and lean mass together, and indicates that the added muscle is functional rather than merely structural.

Muscle mass reflects the balance between protein synthesis and degradation. UCN2 significantly increased myofibrillar fractional synthesis rate in both the sarcopenic obesity and retatrutide cohorts. This occurred without a compensatory rise in protein turnover relative to HFD vehicle, and turnover was lower in UCN2-treated than in retatrutide-treated mice, indicating that muscle accretion is driven primarily by increased synthesis rather than by suppression of degradation. This provides a mechanistic framework for the observed gains in muscle mass and is consistent with previous reports implicating the urocortin/CRHR2 signaling axis in the regulation of skeletal muscle hypertrophy ^18,19^.

The most compelling finding of this study is the ability of UCN2 to preserve skeletal muscle mass and function during incretin-induced weight loss. Lean mass accounts for 25 to 40% of total weight loss on GLP-1R agonists ^2,3,5–8^.Consistent with other observations on semaglutide and tirzepatide ^6,34,38^, retatrutide reduced whole-body lean mass, but TA mass fell by only 9.18%, while liver mass fell by approximately 50%, indicating that a substantial fraction of the apparent lean mass loss is non-muscle tissue. Our data, along with other recently published results ^11^, support that lean mass loss overstates the functional risk, since skeletal force production is preserved in young and old mice. The combination of UCN2 with retatrutide produced the greatest fat mass reduction of any group while preserving TA muscle force and increasing muscle mass. The ability of UCN2 to selectively preserve muscle mass and function during pharmacological weight loss without compromising fat loss offers a potential therapeutic strategy to mitigate these adverse outcomes.

An important consideration in interpreting these results is whether the muscle preserving effects of UCN2 are secondary to reduced food intake or represent a direct action on skeletal muscle. UCN2 treatment reduced food intake in both the sarcopenic obesity and pharmacological weight loss models, consistent with the known anorexigenic effects of urocortin peptides acting through central CRHR2 receptors ^23^. The pair-feeding experiment performed in this study directly addresses this question. and pairfed controls did not show the improvements in TA mass, contractile force, or myofibrillar protein synthesis observed with UCN2 treatment. The energy expenditure data reinforce this dissociation. Pairfed mice showed the expected fall in energy expenditure with caloric restriction, whereas UCN2-treated mice increased expenditure relative to HFD vehicle when analyzed by ANCOVA. UCN2 therefore widens the energy gap by acting on both sides of the balance equation, explaining the greater weight loss achieved relative to matched caloric restriction. Together, these data indicate that the anabolic effects of UCN2 are independent of reduced caloric intake and instead reflect direct action, likely via CRHR2 expressed in skeletal muscle. A consistent feature across cohorts was an immediate shift in substrate selection toward fat oxidation with UCN2 treatment. UCN2 produced a marked fall in RER that was significant within the first 24 h of treatment and persistent throughout the treatment, and was lower than their pairfed counterparts, indicating that the reliance on lipid as fuel is a direct consequence of UCN2 signaling rather than a consequence of decreased food intake. A treatment that simultaneously sustains expenditure and biases oxidation toward lipid oxidation offers a plausible explanation for the fat-selective weight loss and preserved muscle observed with UCN2 treatment. Future work on substrate utilization during UCN2 treatment would be extremely informative to understand this mechanism.

The withdrawal experiment provides perhaps the clearest dissociation between the metabolic and musculoskeletal effects of UCN2. Body weight and fat mass rebounded to near-baseline within 4 weeks, mirroring the relapse after incretin discontinuation, whereas the gain in muscle force conferred by UCN2, alone or with retatrutide, was fully retained, with an overall UCN2 effect on muscle mass of ∼9% by two-way ANOVA. This temporal uncoupling suggests that the anabolic remodeling induced by UCN2 is not merely a passive consequence of the prevailing energy balance but direct adaptations to skeletal muscle.

Questions remain for future work. The downstream signaling pathways mediating the anabolic effects of acylated UCN2 in skeletal muscle require further investigation. This study used only male mice, since females develop obesity more slowly on HFD ^39^; whether UCN2 has comparable efficacy in females, the effects of longer treatment durations, and whether the functional reserve after withdrawal is sustained over longer follow-up all warrant investigation.

In summary, these data demonstrate that a modified long-acting UCN2 preserved skeletal muscle mass and function in models of sarcopenic obesity and pharmacological weight loss, independent of caloric restriction. The combination of UCN2 with retatrutide achieved greater fat loss while increasing muscle force production, and the resulting gains in muscle function persisted for at least 4 weeks after treatment is withdrawn. Together, these data support UCN2 as a therapeutic candidate for sarcopenic obesity and as a combination partner with incretins to improve the composition and durability of pharmacological weight loss.

## Author Contributions

P.V., P.A.M.B., C.D.H. and J.R.G. designed and conducted the studies. J.A.B., P.M.T., J.P.K., C.L.A. and J.T.B. contributed to the design of the studies. E.R.M.Z., D.C.S., W.S.D., A.L.T., E.C.H., E.A., G.S.G., L.P., B.F.M., M.R.W., J.A.W., S.M.B., S.G.L.K. and V.P. conducted experiments. P.V. and P.A.M.B. wrote the manuscript. All authors reviewed and revised the final draft of the manuscript.

## Acknowledgements

We thank the Rodent Metabolic Phenotyping Core (RRID: SCR_022427), supported in part by National Institutes of Health (NIH) grant S10-OD025098, the Cox Institute, and the Institute for Diabetes, Obesity and Metabolism (Perelman School of Medicine, University of Pennsylvania) for performing the semaglutide experiments.

---

**Extended Data Fig. 1.**
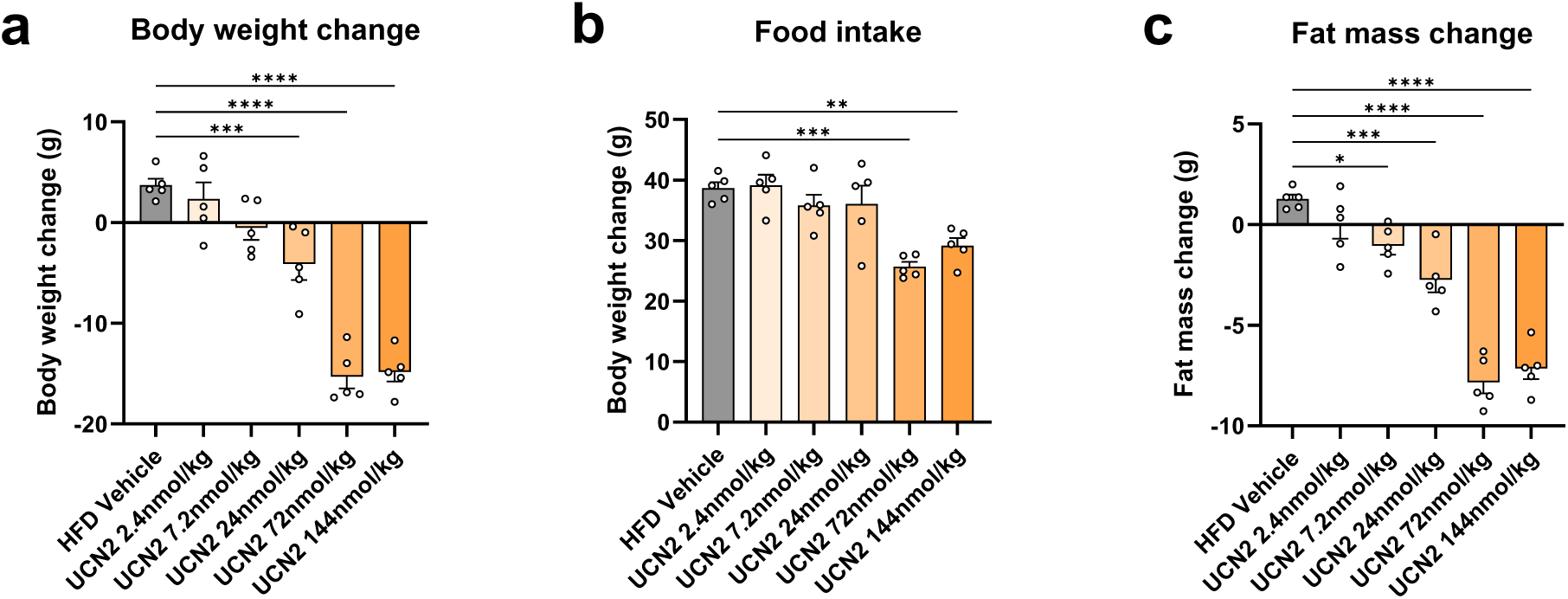
Pharmacologically optimized UCN2 dose-dependently reduces body weight, fat mass and food intake. **a**, Change in body weight in DIO mice treated with vehicle or UCN2 at 2.4, 7.2, 24, 72 or 144 nmol kg⁻ ¹. **b**, Food intake across the same dose range. **c**, Change in fat mass across the same dose range. Data are mean ± s.e.m.; n = 5 mice per group. One-way ANOVA with Tukey’s HSD post hoc test. *P < 0.05, **P < 0.01, ***P < 0.001, ****P < 0.0001.

**Extended Data Fig. 2.**
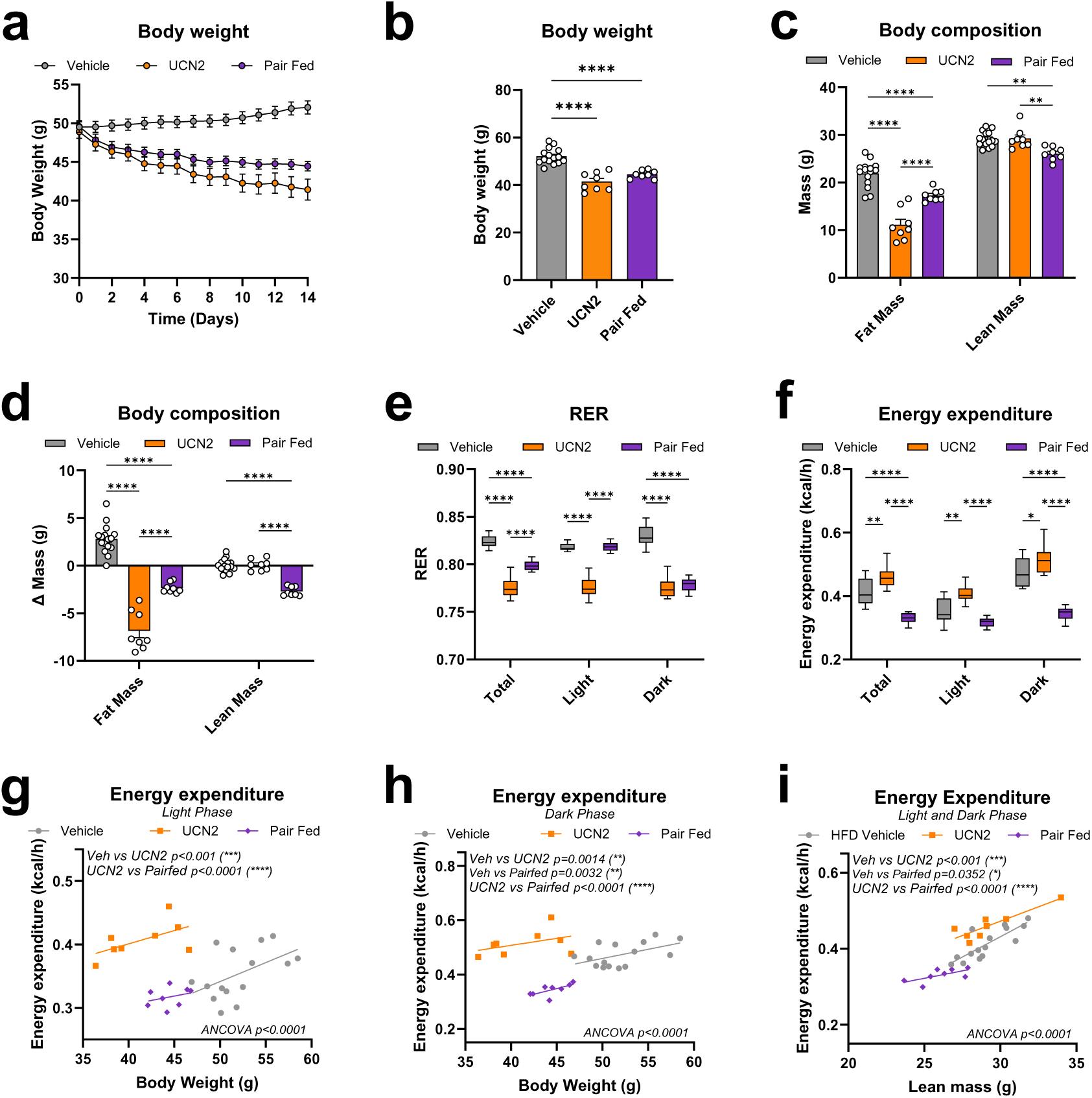
Effects of UCN2 treatment on body composition and energy expenditure at thermoneutrality. **a**, Body weight over the treatment period in mice housed at 30 °C. **b**, Body weight at the end of treatment. **c**, Fat mass and lean mass at the end of treatment. **d**, Change in fat mass and lean mass. **e**, Respiratory exchange ratio (RER) across the total, light and dark phases. **f**, Energy expenditure across the total, light and dark phases. **g**, ANCOVA of light-phase energy expenditure against body weight. **h**, ANCOVA of dark-phase energy expenditure against body weight. **i**, ANCOVA of energy expenditure against lean mass across the light and dark phases. Groups are vehicle, UCN2 and pairfed. Data are mean ± s.e.m.; n = 8-22 mice per group. Energy expenditure was analyzed by ANCOVA with body mass or lean mass as a covariate; remaining comparisons used two-way ANOVA with Tukey’s HSD post hoc test. *P < 0.05, **P < 0.01, ***P < 0.001, ****P < 0.0001.

**Extended Data Fig. 3.**
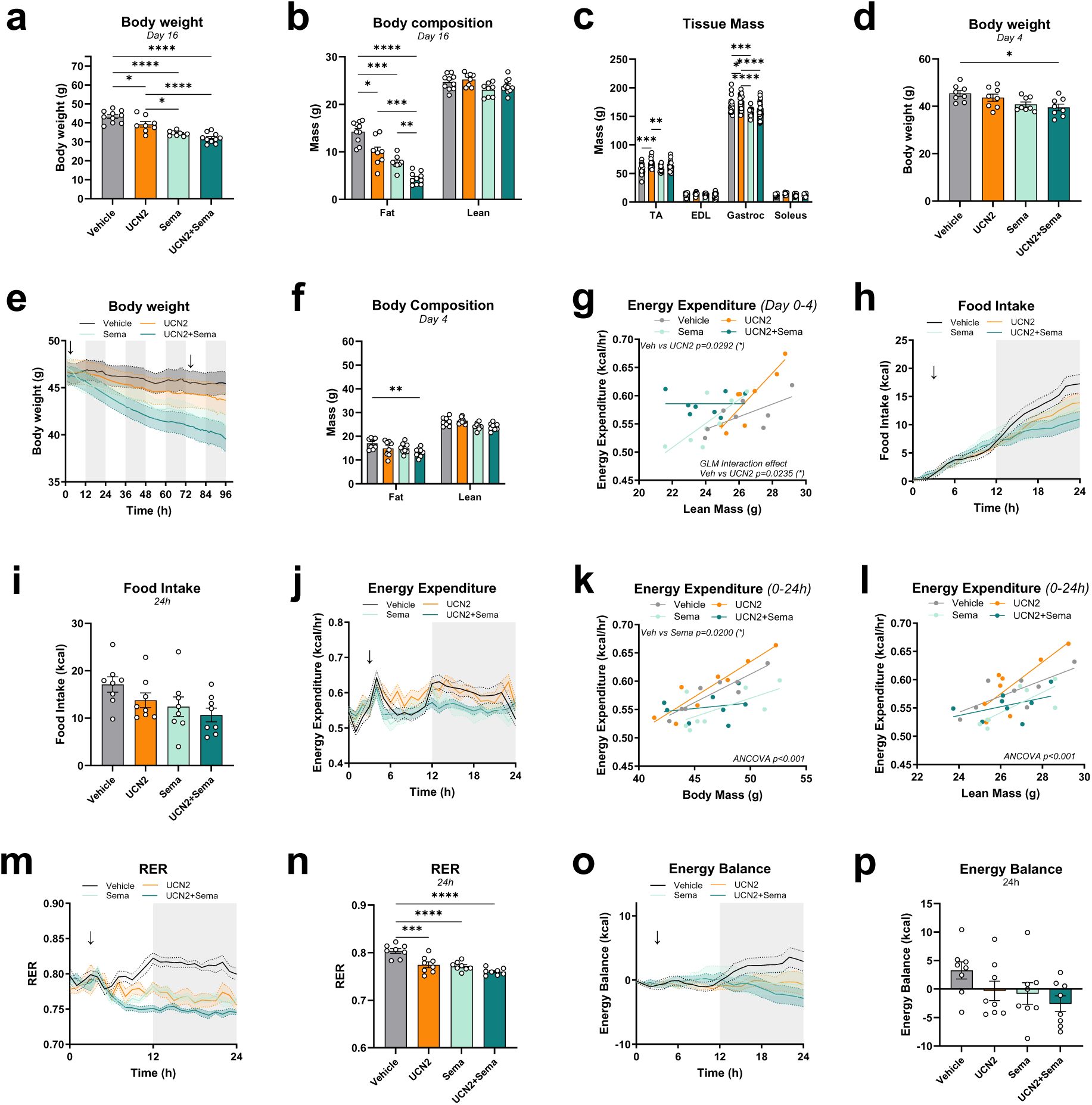
Effects of UCN2 and semaglutide on body composition and energy expenditure. **a**, Body weight at day 16. **b**, Fat mass and lean mass at day 16. **c**, Tibialis anterior (TA), extensor digitorum longus (EDL), gastrocnemius and soleus mass at day 16. **d**, Body weight at day 4. **e**, Body weight over the first 96 h of treatment. **f**, Fat mass and lean mass at day 4. **g**, Average energy expenditure over the first 96h of treatment versus lean body mass with GLM. **h**, Food intake over the first 24 h of treatment. **i**, Cumulative 24 h food intake. **j**, Energy expenditure over the first 24 h of treatment. **k**, Average energy expenditure over the first 24h of treatment versus total body mass with ANCOVA. **l**, Average energy expenditure over the first 24h of treatment versus lean body mass with ANCOVA. **m**, RER over the first 24h of treatment. **n**, Mean 24 h RER. **o**, Energy balance over the first 24 h of treatment. **p**, Cumulative 24 h energy balance. Groups are vehicle, UCN2, semaglutide (sema) and UCN2 + Sema. Data are mean ± s.e.m.; n = 8-10 mice per group. Energy expenditure was analyzed by ANCOVA or a generalized linear model with body mass or lean mass as a covariate; remaining comparisons used one- or two-way ANOVA with Tukey’s HSD post hoc test. *P < 0.05, **P < 0.01, ***P < 0.001, ****P < 0.0001.

**Extended Data Fig. 4.**
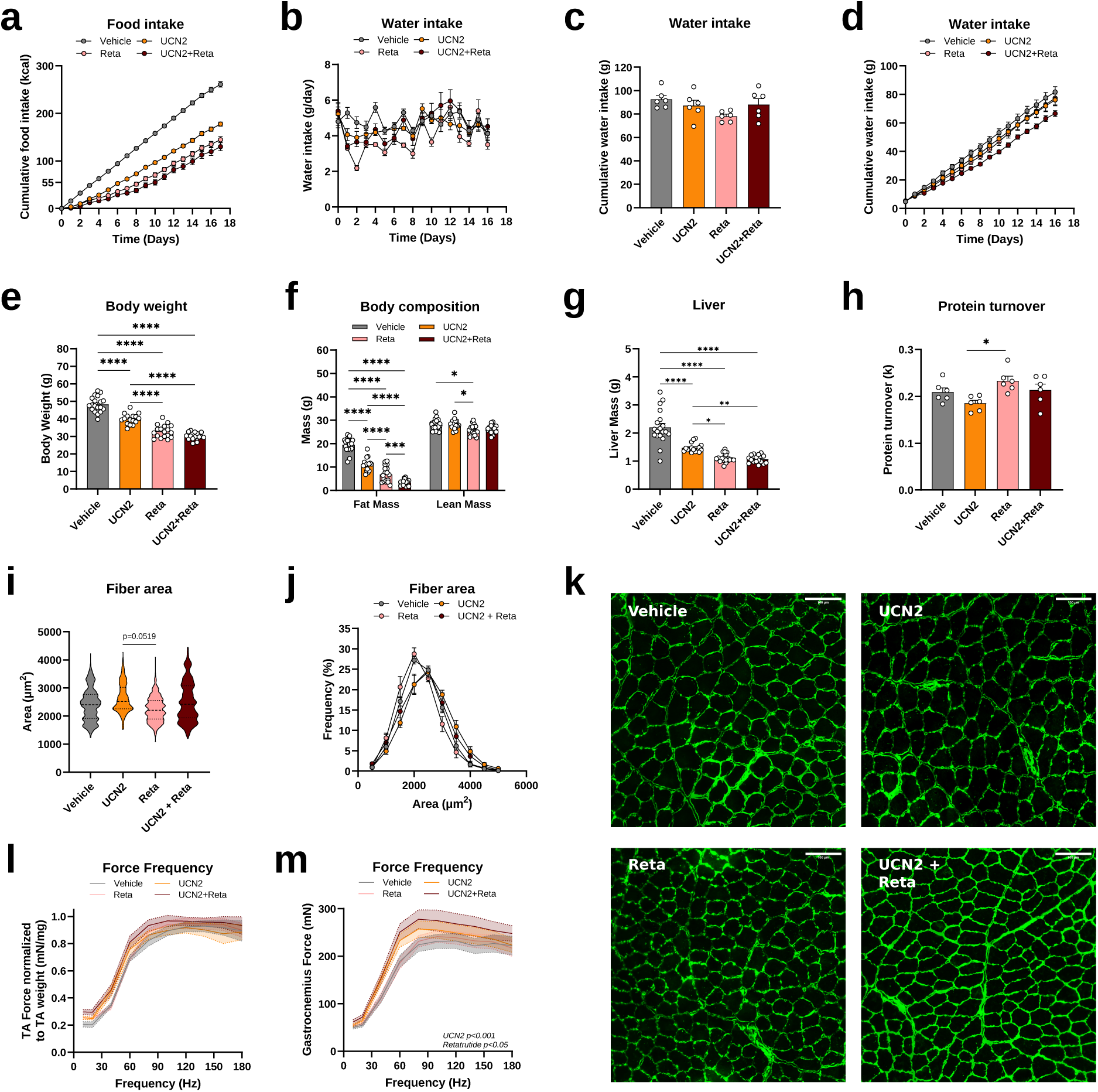
Effects of UCN2 and retatrutide on body composition, muscle mass, force production, protein turnover and liver weight. **a**, Cumulative food intake over the intervention. **b**, Daily water intake. **c**, Cumulative water intake at the end of treatment. **d**, Cumulative water intake over the intervention. **e**, Body weight at the end of treatment. **f**, Fat mass and lean mass at the end of treatment. **g**, Liver mass. **h**, Myofibrillar protein turnover (k). **i**, TA fiber cross-sectional area. **j**, Frequency distribution of TA fiber cross-sectional area. **k**, Representative muscle cross-sections. **l**, TA force–frequency relationship normalized to TA mass. **m**, Gastrocnemius force–frequency relationship. Groups are vehicle, UCN2, retatrutide (reta) and UCN2 + Reta. Data are mean ± s.e.m.; n = 8-22 mice per group. Comparisons used one- or two- way ANOVA with Tukey’s HSD post hoc test. *P < 0.05, **P < 0.01, ***P < 0.001, ****P < 0.0001.

**Extended Data Fig. 5.**
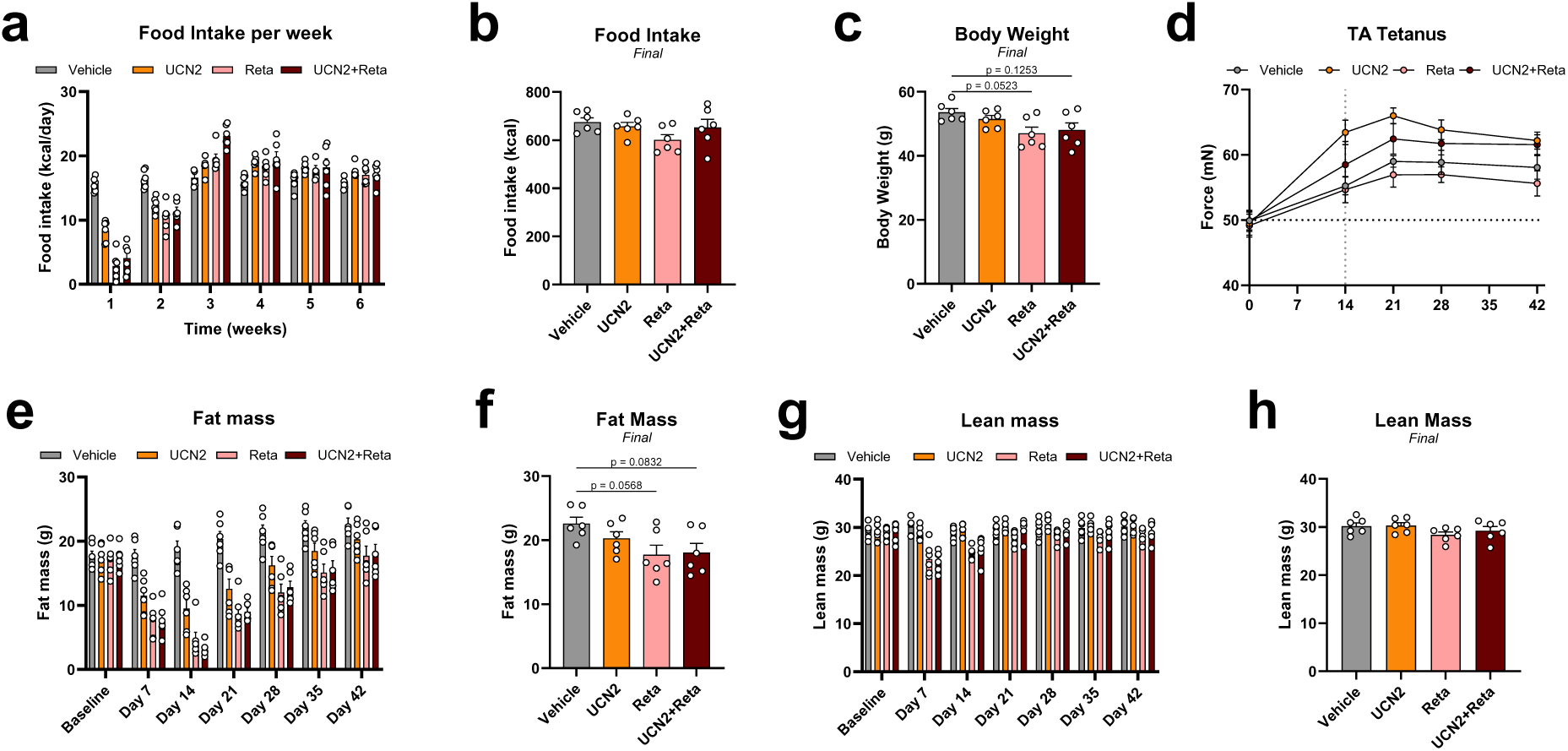
Effects of UCN2 and retatrutide on body composition and force production after cessation of treatment. **a**, Weekly food intake during treatment and withdrawal. **b**, Food intake at the end of the withdrawal period. **c**, Body weight at the end of the withdrawal period. **d**, TA tetanic force during treatment and withdrawal. **e**, Fat mass from baseline through day 42. **f**, Fat mass at the end of the withdrawal period. **g**, Lean mass from baseline through day 42. **h**, Lean mass at the end of the withdrawal period. Mice were treated for 2 weeks and followed for a further 4 weeks after treatment was stopped. Vertical dotted line indicates withdrawal start. Groups are vehicle, UCN2, retatrutide (reta) and UCN2 + Reta. Data are mean ± s.e.m.; n = 6 mice per group. Two-way ANOVA with Tukey’s HSD post hoc test. *P < 0.05, **P < 0.01, ***P < 0.001, ****P < 0.0001.

